# Respiratory tract explant infection dynamics of influenza A virus in California sea lions, northern elephant seals, and rhesus macaques

**DOI:** 10.1101/2020.10.15.342055

**Authors:** Hongwei Liu, Magdalena Plancarte, Erin. E. Ball, Christopher M. Weiss, Omar Gonzales-Viera, Karen Holcomb, Zhong-Min Ma, A. Mark Allen, J. Rachel Reader, Pádraig J. Duignan, Barbie Halaska, Zenab Khan, Divya Kriti, Jayeeta Dutta, Harm van Bakel, Kenneth Jackson, Patricia A. Pesavento, Walter M. Boyce, Lark L. Coffey

## Abstract

To understand susceptibility of wild California sea lions and Northern elephant seals to influenza A virus (IAV), we developed an *ex vivo* respiratory explant model and used it to compare infection kinetics for multiple IAV subtypes. We first established the approach using explants from colonized rhesus macaques, a model for human IAV. Trachea, bronchi, and lungs from 11 California sea lions, 2 Northern elephant seals and 10 rhesus macaques were inoculated within 24 hours post-mortem with 6 strains representing 4 IAV subtypes. Explants from the 3 species showed similar IAV infection kinetics with peak viral titers 48-72 hours post-inoculation that increased by 2-4 log_10_ plaque forming units (PFU)/explant relative to the inoculum. Immunohistochemistry localized IAV infection to apical epithelial cells. These results demonstrate that respiratory tissue explants from wild marine mammals support IAV infection. In the absence of the ability to perform experimental infections of marine mammals, this *ex vivo* culture of respiratory tissues mirrors the *in vivo* environment and serves as a tool to study IAV susceptibility, host-range, and tissue tropism.

**Importance:** Although influenza A virus can infect marine mammals, a dearth of marine mammal cell lines and ethical and logistical challenges prohibiting experimental infections of living marine mammals means that little is known about IAV infection kinetics in these species. We circumvented these limitations by adapting a respiratory tract explant model first to establish the approach with rhesus macaques and then for use with explants from wild marine mammals euthanized for non-respiratory medical conditions. We observed that multiple strains representing 4 IAV subtypes infected trachea, bronchi, and lungs of macaques and marine mammals with variable peak titers and kinetics. This *ex vivo* model can define infection dynamics for IAV in marine mammals. Further, use of explants from animals euthanized for other reasons reduces use of animals in research.

## Introduction

Influenza A viruses (IAV) are important etiologies of respiratory disease in humans and especially affect the elderly, infants, and people with immunodeficiencies and chronic respiratory disease. Dwarfed in 2020 by SARS-CoV-2, IAV are a significant cause of morbidity annually, producing about 500,000 deaths worldwide each year (1). IAV possess a wide host range that includes birds, horses, pigs, and humans (2, 3). Marine mammals can also be infected, sometimes with strains from human pandemics (4, 5). The viral genetic and host factors that affect cross-species transmission by IAV, especially from birds to mammals including humans, have been extensively studied (5–9). However, mechanisms of zoonotic IAV transmission from avian or human to other mammalian species, including marine mammals, are less well understood (6).

IAV was first identified in North American marine mammals in 1979, when an H7N7 epizootic killed 500 harbor seals and caused hemorrhagic pneumonia in others (10–12). Since then, IAV infection and sometimes disease has been identified in several marine mammal species, including mass mortalities in harbor seals near Cape Cod, Massachusetts, USA, attributed to H7N7, H4N5, or H4N6, and detection of antibody to multiple H and N subtypes in several seal species (13–20). On the West Coast, surveillance from 2009 to 2015 (4, 21, 22), and from 2016 to present (unpublished) in multiple marine mammal species from California shows variable IAV exposure. Seroprevalence in sea otters and Northern elephant seals is higher than in sympatric harbor seals and California sea lions, and sea otters are exposed to avian- and human-origin IAV, including pandemic H1N1 (4, 21). Despite frequent antibody detection, isolation of IAV from marine mammals has been limited. In California, only pandemic H1N1 has been isolated from 2 Northern elephant seals in 2010 (4). Together, these surveillance data suggest that marine mammals can be infected with some of the same IAV subtypes that cause human epidemics. However, in the absence of *in vivo* study capabilities in marine mammals, infection dynamics and subtype-specific susceptibility remain unknown. Therefore, there is a need to develop approaches to study the infection biology of IAV in these species.

Given that the respiratory tract is the initial site of IAV infection, defining infection dynamics and cellular tropism in respiratory tissues is important for assessing species susceptibility. However, *in vivo* systems for studying susceptibility of marine mammals to IAV are not available. As an alternative, *ex vivo* explants from respiratory tract tissues can mimic the physiological microenvironment of a respiratory tract. *Ex vivo* systems in human and animals (excluding marine mammals) have been successfully developed and used to study host innate responses, infection dynamics, viral genetic determinants of infection, antiviral drug treatments, and pathogenesis of human and animal IAV and other respiratory viruses (23–28). The goal of this study, therefore, was to expand these existing models for use in marine mammals to study susceptibility and infection dynamics of IAV strains of mammalian and avian origin to assess the potential for interspecies transmission, and to providing a new approach for studying the biology and pathogenesis of IAV in marine mammals. Given that marine mammal tissues are only opportunistically available from wild animals treated at The California Marine Mammal Center in Sausalito, CA, USA, we first established the *ex vivo* explant system using rhesus macaques that are more regularly available from the California National Primate Research Center, Davis, CA, USA. Rhesus macaques represent a valuable model for understanding IAV infection dynamics in the human respiratory tract due to similar structure, physiology, and mucosal immunity. Additionally, IAV infection dynamics have not been studied in rhesus macaques, although they are a model for human IAV infection (29) and are used for testing vaccine candidates (30, 31).

We used trachea, bronchi, and lung explants from rhesus macaques to compare the susceptibility, infection kinetics, and tissue tropism of 6 strains of IAV from 4 subtypes to validate the utility of this system. After establishing the explant approach with IAV in rhesus macaques, we used it to study IAV infection dynamics and tropism in California sea lions and Northern elephant seals. We observed that both rhesus macaques and marine mammals are susceptible to all 6 IAV strains, and that rhesus macaque and California sea lion respiratory tract explants exhibit temporal, tissue, and IAV subtype-dependent IAV infection kinetics.

## Material and Methods

### Ethics Statement

This project was conducted with approval from the United States National Marine Fisheries Service Marine Mammal permit # 18786-04. The University of California, Davis is accredited by the Association for Assessment and Accreditation Laboratory Animal Care International (AAALAC). Animal care was performed in compliance with the 2011 Guide for the Care and Use of Laboratory Animals provided by the Institute for Laboratory Animal Research. Rhesus macaque studies were approved by the University of California, Davis IACUC under protocol #19760.

#### Cell culture

Madin-Darby Canine Kidney (MDCK) cells (ATCC #CCL-34) were maintained at 37°C and 5% CO_2_ in MDCK-growth-medium (Iscove’s Modified Dulbecco’s Medium (IMDM), supplemented with 5% FBS, 0.1% sodium bicarbonate, penicillin and streptomycin).

#### Influenza A virus propagation and titration

We used IAV strains of avian and marine mammal origin for explant studies (**Table 1**). The harbor seal H3N8 (HS/H3N8) IAV strain was obtained from Dr. Hon Ip, (National Wildlife Health Center, Madison Wisconsin). The other strains were isolated at UC Davis and used in this study. Avian allantoic fluid virus stocks of 6 strains were amplified in Madin-Darby canine kidney (MDCK) cells to obtain sufficient titers and volumes for use as inocula. Viral titers of the 6 inocula were determined by plaque assay using MDCK cells. Briefly, MDCK cells were grown in 6-well plates to 80-90% confluence in MDCK growth medium. Viral stocks were serially ten-fold diluted in viral-growth-medium (Minimum Essential Medium (MEM), supplemented with 0.5% BSA, 0.1% sodium bicarbonate, 10 mM Hepes, penicillin and streptomycin). The MDCK cells were washed 3 times with Dulbecco’s phosphate buffered saline (DPBS) prior to inoculation with 200 ul of diluted virus samples. After 1 hour incubation at 37°C with 5% CO_2_, 3 ml viral-growth-medium containing 1 ug/ml N-p-tosyl-L-phenylalaninechloromethyl ketone-treated (TPCK) and 0.5% agarose was added to each well. After a 48 hour incubation at 37°C with 5% CO_2_, MDCK cells were fixed with 4% formaldehyde and stained with 0.05% crystal violet.

**Table 1.** IAV strains used for explant infection experiments. ES is elephant seal, HS is harbor seal, m is mallard. N/A indicates strain was isolated at UC Davis and first described in this study. MDCK is Madin-Darby canine kidney cell. PFU is plaque forming unit.

| **Virus ID** | **Strain name** | **Subtype** | **Passage number** | **MDCK or Vero E6 titer (PFU/ml)** | **GenBank Accession number** | **Reference** |
| --- | --- | --- | --- | --- | --- | --- |
| ES/H1N1 | A/Elephant seal/California/1/2010 | H1N1 | p4 | $1.3 \times 10^8$ | JX865419-JX865426 | Goldstein et al. 2013 (4) |
| HS/H3N8 | A/harbor seal/New Hampshire/179629/2011 | H3N8 | p3 | $1.8 \times 10^6$ | KJ467564-KJ467571 | Karlsson et al. 2014 (5) |
| m/H3N8 | A/mallard/California/1475/2010 | H3N8 | p2 | $6 \times 10^6$ | CY120501-CY120508 | N/A |
| m/H1N1 | A/mallard/California/3134/2010 | H1N1 | p2 | $4 \times 10^7$ | CY120611-CY120618 | N/A |
| m/H5N2 | A/mallard/California/2396/2010 | H5N2 | p2 | $4 \times 10^7$ | CY120587-CY120594 | N/A |
| m/H7N5 | A/mallard/California/1390/2010 | H7N5 | p2 | $1 \times 10^7$ | CY120555-CY120562 | N/A |

#### In vitro infection kinetics of IAV

Infection kinetics for the 6 IAV strains were assessed in MDCK cells. Cells were grown at 37°C with 5% CO_2_ to 6 × 10^5^ cells/well in 24-well plates and inoculated in triplicate with 6 × 10^3^ PFU virus of IAV, representing a multiplicity of infection (MOI) of 0.01. After 1 hour incubation, each well was washed with 1 ml of DPBS 3 times to remove unbound virus. Next, viral growth medium supplemented with 1 ug/ml TPCK trypsin for IAV was added to each well. The supernatant was sampled at 1, 24, 48, and 72 hours post-inoculation and stored at −80°C in virus growth medium. Viral titers were assessed by plaque assay.

#### Titration to quantify infectious IAV

Plaques were counted against a white background and the titer in plaque forming units (PFU) was determined by counting wells with individual plaques. Two to 3 dilutions of each explant were tested once. Viral titers were recorded as the reciprocal of the highest dilution where plaques are noted and represented as PFU per ml for liquid samples or PFU per explant for tissues. The limit of detection (LOD) of the plaque assay was 50 PFU/ml or explant.

#### Explant sources, collection and processing

Respiratory tract tissues including trachea, bronchi, and lungs were used for explant studies (**Table 2**). Tissues were obtained from Indian origin rhesus macaques (*Macaca mulatta*), euthanized due to medical conditions, who were born and raised at the California National Primate Research Center, University of California, Davis, CA. The respiratory tract tissues of marine mammals were from wild California sea lions (*Zalophus californianus*) and Northern elephant seals (*Mirounga angustirostris*) who stranded on beaches and were euthanized due to medical conditions at The Marine Mammal Center in Sausalito, CA. The Marine Mammal Center rehabilitates stranded marine mammals. Some animals that present with severe conditions are not able to be returned to the wild and are therefore euthanized. To reduce confounding effects of respiratory disease on IAV infection, we intentionally excluded animals whose necropsy reports indicated respiratory disease as the primary or sole reason for euthanasia. Serum harvested from blood collected at necropsy from macaques and marine mammals were tested for IAV antibody directed against a conserved epitope of the nucleoprotein using an enzyme-linked immunosorbent assay (ELISA) ID-Screen® Influenza A Antibody Competition Multi-species kit, (IDvet, Grabels, France) following the manufacturer’s instructions. Positive and negative controls included in the kit were also tested on each plate. An ELX808 BioTek Spectrophotometer (BioTek Instruments, Winooski, VT) was used to measure absorbance. Sera was considered positive for IAV antibody when the ratio of the absorbance of the test sample to the negative control was less than 0.45, as previously established (21, 22). Marine mammals were also tested for IAV RNA by reverse transcription polymerase chain reaction (RT-PCR). Nasal and rectal swabs were collected from each California sea lion and Northern elephant seal by veterinary staff at The Marine Mammal Center upon entry to the facility. Swabs were placed in vials containing 1.5 ml of viral transport media (VTM). Samples were refrigerated for up to one week prior to shipping to the laboratory. Once received at the laboratory, swab samples were processed the same day or stored at −80°C. RNA was extracted from swab samples using the MagMAX-96 AI/ND Viral RNA Isolation Kit (Applied Biosystems, Foster City, CA) and a KingFisher Magnetic Particle Processor (Thermo Scientific, Waltham, MA). Extracted RNAs from swab samples were subjected to an established IAV RT-PCR (32) targeting a conserved region of the matrix gene using the AgPath-IDTM One Step RT-PCR mix (Applied Biosystems, Foster City, CA) and an ABI 7500 real-time PCR System (Applied Biosystems, Foster City, CA). The positive control, a cell culture isolate of ES/H1N1, and a negative control, VTM, were also tested on each plate. Samples with a cycle threshold (Ct) value <45 were considered positive (33). Tissues from all three species were stored for up to 24 hours in Roswell Park Memorial Institute-1640 (RPMI) medium (Gibco) at 4°C. During tissue preparations, lungworms (*Parafilaroides decorus*) were observed, either in airways (i.e., bronchioles/bronchi) or sometimes free in the lung parenchyma (in which case they were occasionally associated with mild inflammation), in all California sea lions. Areas of tissues without worms were selected as explants. No worms were observed grossly in explants during the IAV infection studies. Prior to preparation for infection studies, tissues were separated and washed 4-6 times in 50 ml conical tubes containing 30 ml of RPMI medium for 2-5 min each at 27°C. The final wash was performed in RPMI-growth-medium (RPMI medium supplemented with 100 U/ml each of penicillin and 100 ug/ml streptomycin (Gibco) and 1x antibiotic-antimycotic (Gibco)).

**Table 2.**
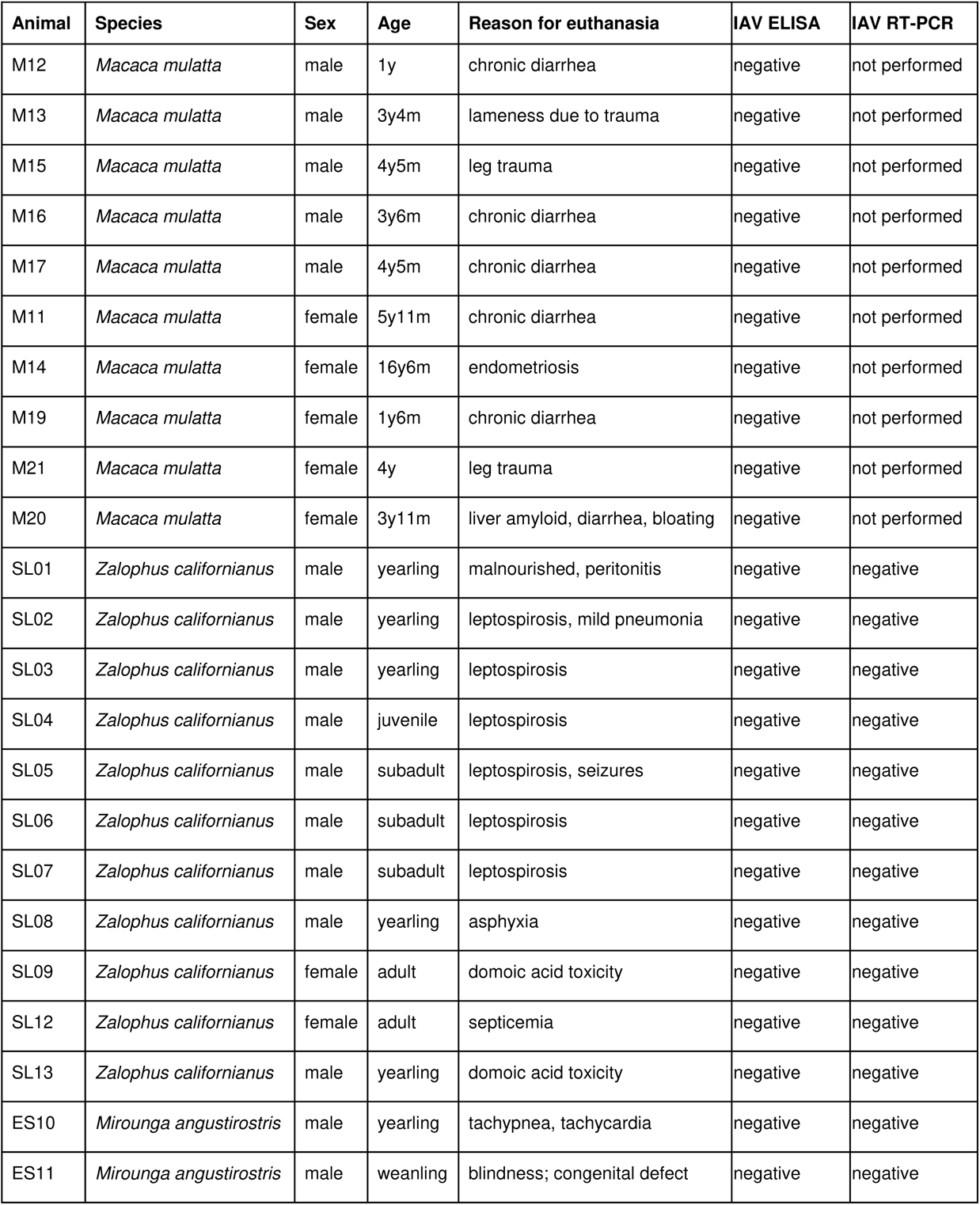
Explant sources for IAV *ex vivo* infection studies. Tissues were derived from 10 rhesus macaques (*Macaca mulatta*), 11 California sea lions (*Zalophus californianus*) and 2 northern elephant seals (*Mirounga angustirostris)*. M is macaque, SL is sea lion, ES is elephant seal, y is year, m is month.

| Animal | Species | Sex | Age | Reason for euthanasia | IAV ELISA | IAV RT-PCR |
| --- | --- | --- | --- | --- | --- | --- |
| M12 | *Macaca mulatta* | male | 1y | chronic diarrhea | negative | not performed |
| M13 | *Macaca mulatta* | male | 3y4m | lameness due to trauma | negative | not performed |
| M15 | *Macaca mulatta* | male | 4y5m | leg trauma | negative | not performed |
| M16 | *Macaca mulatta* | male | 3y6m | chronic diarrhea | negative | not performed |
| M17 | *Macaca mulatta* | male | 4y5m | chronic diarrhea | negative | not performed |
| M11 | *Macaca mulatta* | female | 5y11m | chronic diarrhea | negative | not performed |
| M14 | *Macaca mulatta* | female | 16y6m | endometriosis | negative | not performed |
| M19 | *Macaca mulatta* | female | 1y6m | chronic diarrhea | negative | not performed |
| M21 | *Macaca mulatta* | female | 4y | leg trauma | negative | not performed |
| M20 | *Macaca mulatta* | female | 3y11m | liver amyloid, diarrhea, bloating | negative | not performed |
| SL01 | *Zalophus californianus* | male | yearling | malnourished, peritonitis | negative | negative |
| SL02 | *Zalophus californianus* | male | yearling | leptospirosis, mild pneumonia | negative | negative |
| SL03 | *Zalophus californianus* | male | yearling | leptospirosis | negative | negative |
| SL04 | *Zalophus californianus* | male | juvenile | leptospirosis | negative | negative |
| SL05 | *Zalophus californianus* | male | subadult | leptospirosis, seizures | negative | negative |
| SL06 | *Zalophus californianus* | male | subadult | leptospirosis | negative | negative |
| SL07 | *Zalophus californianus* | male | subadult | leptospirosis | negative | negative |
| SL08 | *Zalophus californianus* | male | yearling | asphyxia | negative | negative |
| SL09 | *Zalophus californianus* | female | adult | domoic acid toxicity | negative | negative |
| SL12 | *Zalophus californianus* | female | adult | septicemia | negative | negative |
| SL13 | *Zalophus californianus* | male | yearling | domoic acid toxicity | negative | negative |
| ES10 | *Mirounga angustirostris* | male | yearling | tachypnea, tachycardia | negative | negative |
| ES11 | *Mirounga angustirostris* | male | weanling | blindness; congenital defect | negative | negative |

#### Preparation of tracheal and bronchi explants

A simplified *ex vivo* culture procedure was derived from the methods described by Nunes *et al*. (Nunes et al, 2010). After washing in RPMI medium, the surrounding connective tissue exterior to the tracheal and bronchi cartilage was removed. Trachea were cut into O-rings horizontally where each slice contained about 0.5 cm cartilage. Each O-ring consisted of the respiratory mucosa, epithelial cell layer, and underlying cartilage. O-rings were further cut into small pieces of approximately 0.5 cm × 0.5 cm. Bronchi were cut similarly. Explants were implanted singly onto agarose plugs (RPMI-growth-medium containing 0.5 % agarose) in 24-well plates with the mucosal surface facing up. Explants were maintained in a humidified 37°C incubator with 5% CO_2_ for up to 7 days.

#### Preparation of lung, liver, and kidney explants

Lung tissues were used to investigate IAV tissue specificity. The liver and kidney of two rhesus macaques were also used as non-target tissue controls where IAV infection was not expected. Processing of these tissues was the same as for trachea and bronchi, where tissues were cut into pieces of approximately 0.5 cm^3^ after washing. Each piece was implanted onto agarose plugs and incubated using the same conditions as for trachea and bronchi.

#### Ex vivo IAV infection kinetics

To evaluate IAV infection kinetics *ex vivo*, trachea, bronchi, and lung explants were inoculated in triplicate (when sufficient tissue was available) with 1 × 10^4^ PFU of one of each of the 6 IAV strains in 10 ul viral growth medium onto the epithelial surface of each section for rhesus macaques. The inoculum volume for marine mammals was adjusted to 20 ul at a dose of 2 × 10^4^ PFU for each IAV strain. Viral growth medium was used in the mock-infected control explants. Inoculated explants were sampled at 1 hour post inoculation (hpi) and every 24 hours up to 72 hpi (or as indicated in graphs) for viral quantification and histology. For viral quantification, each infected explant was directly immersed in 0.5 ml RPMI-growth-medium in a 2 ml centrifuge tube with a 5 mm glass bead and homogenized in a Mixer Mill MM300 (Retsch, Leeds, UK) at 30 hertz for 4 minutes at 22°C followed by centrifugation at 16,000 g for 4 minutes and storage at −80°C. Titers of the released progeny virus from explants were determined by plaque assay. Titers are reported as the geometric mean of triplicate explants at each time point.

#### Stability assays

To determine whether IAV detected after inoculation was due to productive virus infection in respiratory tissues, we performed viral stability assays in the absence of tissue sections. Medium containing 1 × 10^4^ PFU/ml of each of the 6 IAV strains was added to 24-well tissue culture plates and supernatant was collected at 0, 24 and 48 hpi for the determination of viral titers by plaque assay. As a second step, the stability assays were also performed using non-target tissues (liver and kidney) to confirm that the viral titers detected in respiratory tissues reflect productive infection in target cells. Liver and kidney explants were inoculated with 1 × 10^4^ PFU of each of the 6 IAV strains in 10 ul RPMI growth medium and assayed by the same methods used for respiratory tract tissues.

#### IAV immunohistochemistry

Tracheal, bronchial, and lung explants from a subset of IAV-inoculated animals (M11, M13, SL08, and SL12) were fixed in 10% buffered formalin and embedded in paraffin. Explants not treated with IAV and incubated only with rabbit IgG (Invitrogen) as primary antibody were included as negative isotype controls for background staining. These animals were selected since they showed higher IAV titers than others. Antigen retrieval was performed on 5 um sections via incubation in AR10 (Biogenex) in a digital decloaking chamber (Biocare Medical) for 2 minutes at 125°C, followed by cooling to 90°C for 10 minutes, rinsing with water, and a final Tris-buffered saline with 0.05% Tween 20 rinse. Tissue sections were exposed to the primary rabbit anti-influenza A virus nucleoprotein antibody (LSBio), at a ratio of 1:250 with antibody diluent (Dako). Tris-buffered saline with 0.05% Tween 20 was used for all washes. Nonspecific binding sites were blocked with 5% bovine serum albumin (Jackson ImmunoResearch). Binding of the primary antibody was detected using Envision rabbit polymer with AEC as the chromogen (Dako). Each tissue section was evaluated independently by two pathologists. Sections from slides were visualized with a Zeiss Imager Z1 (Carl Zeiss). Digital images were captured and analyzed using Openlab software (Improvisation). Cells with nuclear immunoreactivity were considered positive. The positive cells in the epithelium of the trachea were manually counted. The area of the tracheal epithelium was measured. The number of positive cells was presented as cells per square millimeter of the epithelium.

#### Influenza A virus genome sequencing

IAV RNA from inocula and homogenized explant samples harvested 48 to 96 hpi from selected animals for all 3 species were sequenced. Viral RNAs were extracted using the Magmax-96 AI/ND Viral RNA isolation kit (Applied Biosystems). Viral RNA extracts were used as a template for multi-segment RT-PCR reactions to generate all 8 genomic segments of IAV using procedures described previously (22, 34). Consensus sequences were generated at the Icahn School of Medicine at Mount Sinai, as described previously (35). Genome sequences from explant samples were compared to inocula.

#### Statistical analyses

Statistical analyses were conducted using GraphPad Prism software version 8. Two-way ANOVA with Dunnett’s multiple comparisons tests were performed to compare the mean viral titers in explants at 24, 48 and 72 hpi to the mean titer 1 hpi as well as mean viral titers in MDCK cells for all 6 IAV strains at 24, 48 and 72 hpi. IAV titers in tissue explants were plotted at 0, 24, 48, and 72 hpi and mean area under the curve (AUC) was calculated for all titers above the assay limit of detection. Within individual tissues, mean AUC was compared by one-way ANOVA using Tukey’s method for multiple comparisons between all IAV strains. AUC for California sea lions grouped by reason for euthanasia (respiratory versus non-respiratory) were compared by Mann-Whitney rank test. Mean titers for IAV strains and tissues after inoculation of rhesus macaque and California sea lion explants were computed using R (36). Welch ANOVA with the Games-Howell post-hoc tests were used to compare changes in mean log_10_ viral titers from 1 to 24, 48, and 72 hpi across IAV strains, tissues within species, and by IAV strain and tissue pairs between species.

#### Data availability

Sequencing data are available at the CEIRS Data Processing and Coordinating Center (DPCC) Project Identifier: SP4-Boyce_5004 and Submission_ID: 1136547734004 and at GenBank accession numbers MW132167-MW132399.

## Results

### Influenza A virus strain selection

We compared relative infection kinetics for 6 IAV strains (**Table 1**) used in this study. Since marine mammals and water birds share the same near shore environments where cross-species transmission can occur, we used strains isolated from both marine mammals and birds. The strains used also represent common IAV subtypes. Further, the elephant seal (ES)/H1N1 strain shows high genetic similarity with human pandemic H1N1 (4). Strains were isolated from an elephant seal (ES, N=1), or a harbor seal (HS, N=1), or mallard ducks (m, N=4), and they share similar passage histories in embryonated chicken eggs and MDCK cells.

### Influenza A virus strains exhibit differential infection kinetics in immortalized cells

Given that cell infection kinetics for 4 of the strains had not previously been established, we first performed growth assays for each of the 6 IAV strains after inoculation into MDCK cells in triplicate at a MOI of 0.01 (**Figure 1**). The H3N8 strain isolated from a harbor seal (HS/H3N8), exhibited the slowest and lowest growth kinetics, peaking at 10^5^ PFU/ml 48 hpi. The H1N1 strains showed the fastest and highest growth kinetics, peaking at >10^7^ PFU/ml by 24 hpi. The viral infection kinetics of the other 4 strains tested, 3 of which were isolated from mallard ducks and 1 from a harbor seal, were intermediate between HS/H3N8 and the 2 H1N1 strains. Titers for all strains were significantly higher than those for HS/H3N8 at one or more time points between 24-72 hpi (two way ANOVA). Given these data show that *in vitro* infection kinetics in an immortalized cell line are different for the 6 IAV strains used in this study, we sought to determine whether the growth kinetics of these strains also differed in explant tissues from the 3 animal species.

**Figure 1.**
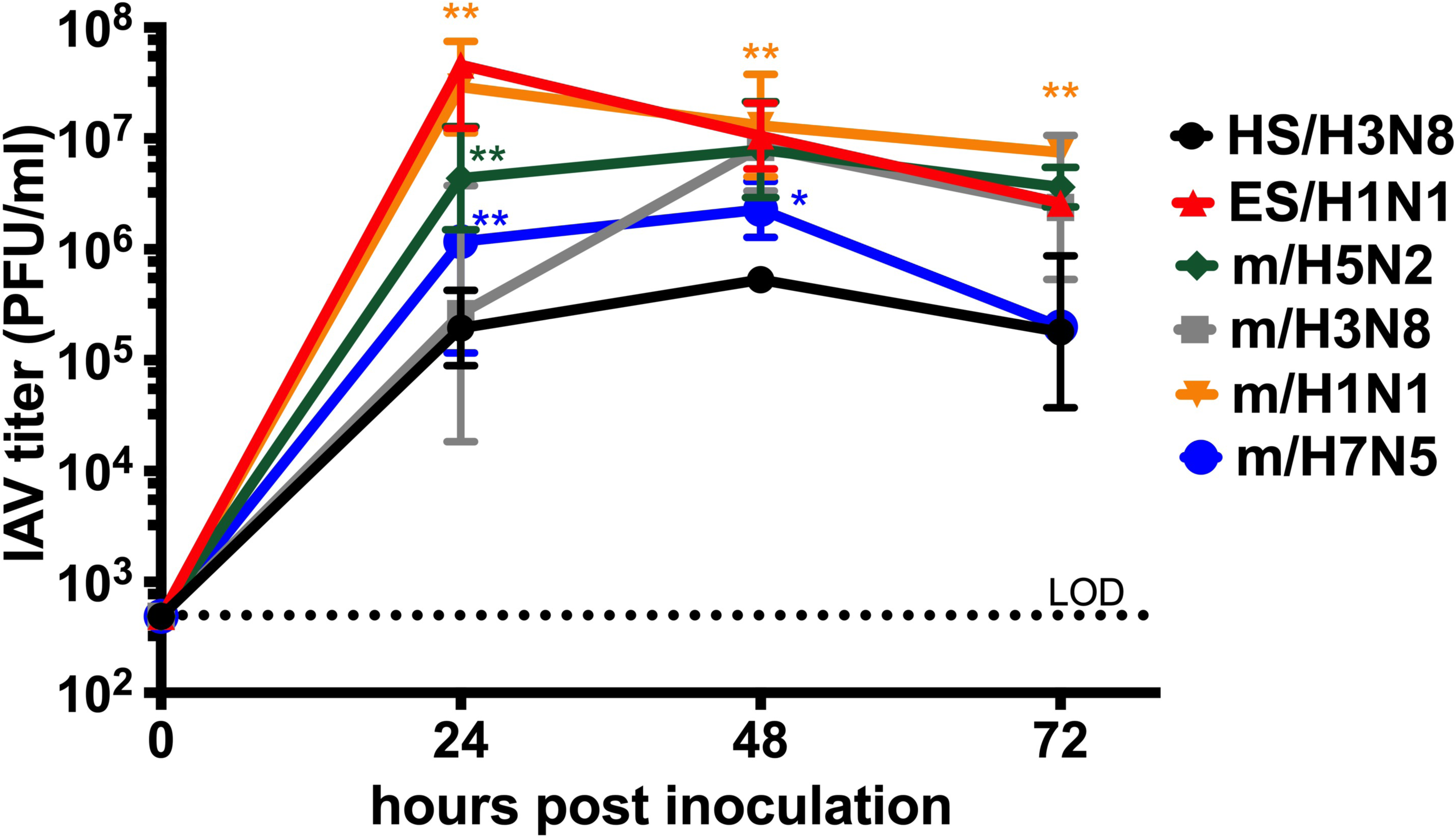
*In vitro* infection kinetics of 6 influenza A virus strains in Madin-Darby canine kidney cells. A MOI of 0.01 was used. The geometric mean virus titer from triplicate wells at each time point was plotted as PFU per milliliter ± geometric standard deviation. The dotted line represents the limit of detection (LOD) of 500 PFU/ml. Asterisks show comparisons of mean titers between HS/H3N8 and all other strains analyzed by two-way ANOVA of log transformed values where * is *p* < 0.001 and ** is *p* < 0.0001. Strain names refer to the original source of the virus isolate followed by the hemagglutinin and neuraminidase subtype. HS is harbor seal. m is mallard. ES is elephant seal.

### Influenza A viruses infect rhesus macaque respiratory tract explants

Given that marine mammal tissues are only opportunistically available from wild animals treated at The California Marine Mammal Center in Sausalito, CA, USA, we first established the *ex vivo* explant system using rhesus macaques that are more regularly available from the California National Primate Research Center, Davis, CA, USA (**Figure 2A**). The rhesus macaques were bred and then euthanized at the California National Primate Research Center due to non-respiratory medical conditions. Although tissues were used within 6 h post-mortem, we microscopically visualized movement of glass beads by cilia placed atop tracheas from some macaques and verified viability at 24, 48, and 72 hours, similar to Nunes *et al*. (23). To determine whether rhesus macaque explants support IAV infection and to define infection kinetics, we measured viral titers for 7 days (from 1 to 168 hpi) in explants from the first animal, an 11-year old female macaque inoculated with HS/H3N8 (**Figure 2B**) who had no detectable IAV antibody in sera at the time of necropsy. We defined infection as detection of a kinetic increase in infectious IAV titer above the inoculum, 10^4^ PFU. Titers increased over time in all 3 tissues and exceeded inocula from 48-168 hpi in trachea, from 24-120 hpi in bronchi, and from 72-96 hpi in the lung. Maximal titers >10^5^ PFU/tissue were detected in all 3 tissue types. Decreases in titers over time (i.e. 120-168 hpi) were likely concomitant with death of target cells in explants. By contrast, when 10^4^ PFU of IAV strains were inoculated into medium in the absence of explants (**Figure 2C**) or into non-IAV target kidney (**Figure 2D**) or liver (**Figure 2E**) from a 3.5 year old macaque, no IAV was detected above the limit of detection (50 PFU/explant) at 24 or 48 hpi, which indicates IAV are not viable over time absent IAV-susceptible tissues. Together these results demonstrate that rhesus macaque respiratory tract explants support productive infection by IAV.

**Figure 2.**
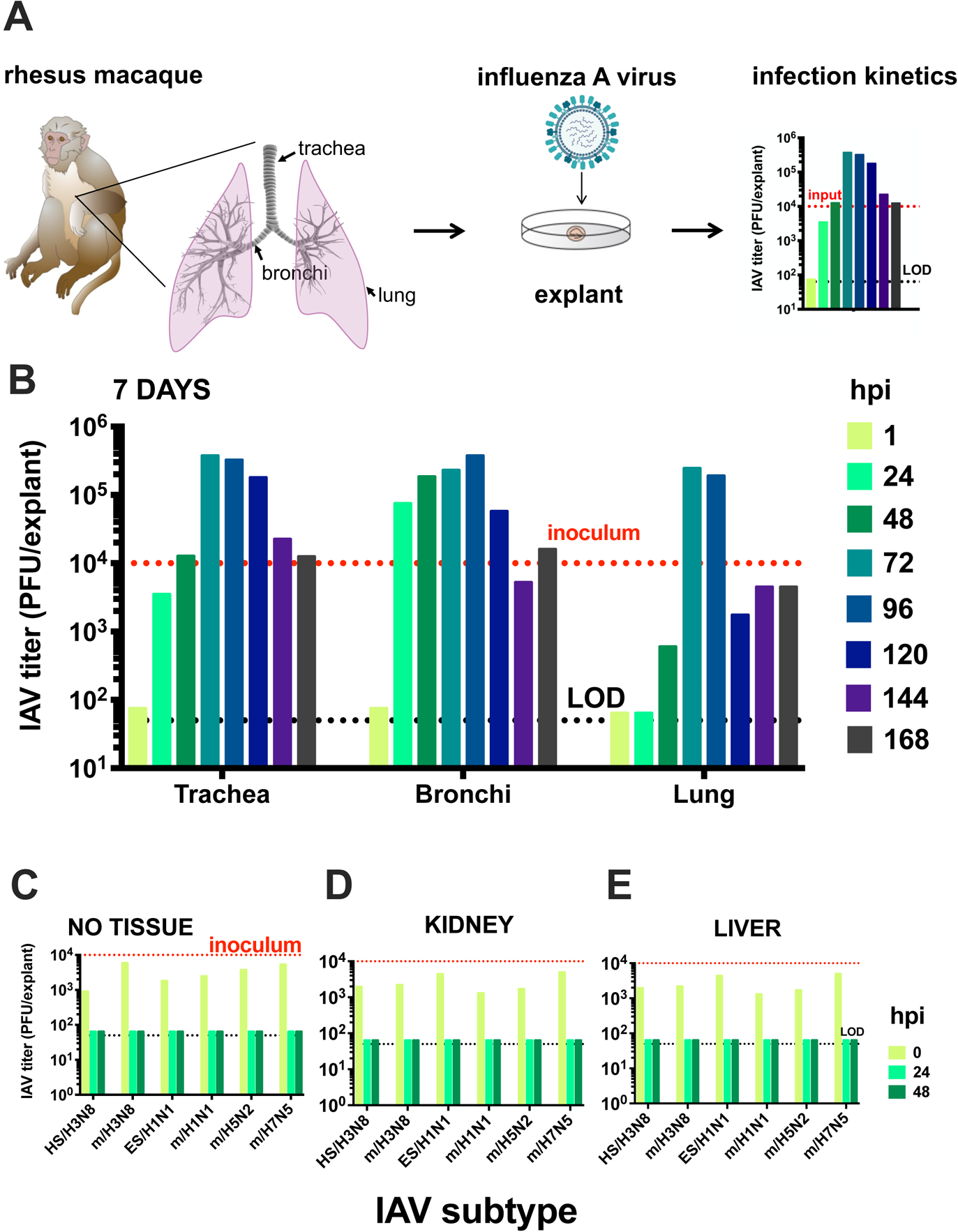
**(A) Experimental design for *ex vivo* rhesus macaque respiratory tract explant inoculations with influenza A virus (IAV). (B)** Seven day infection kinetics of IAV HS/H3N8 in explants from an 11 year old animal that was serologically IAV non-reactive at necropsy. **(C)** IAV titers in the absence of explants to evaluate stability of infectious virus from 0 to 48 hpi. IAV titers in kidney **(D)** and liver **(E)**, tissues that are not IAV targets from a 3.5 year old animal from 0 to 48 hpi. Each bar shows the measurement from a single explant. The dotted black line shows the limit of detection, 50 PFU/explant. The red line shows the inoculum. hpi is hours post inoculation. Strain names refer to the original source of the virus isolate followed by the hemagglutinin and neuraminidase subtype. HS is harbor seal. m is mallard. ES is elephant seal.

### Influenza A virus exhibits strain-specific infection patterns in ex vivo respiratory tract tissues

#### Rhesus macaques

To determine whether rhesus macaques share similar susceptibilities to different IAV subtypes, we inoculated respiratory tract explants from 10 rhesus macaques (**Table 2**) with 6 virus strains. Five of the macaques were females and five were males and they ranged from 1 to 16 years of age. Most of the macaques were euthanized due to injuries sustained due to non-respiratory conditions including trauma or chronic diarrhea. All macaques tested IAV seronegative by ELISA in serum from blood collected at necropsy (**Table 2**). We focused on 0 to 72 hpi since increases in IAV titers were observed over that period in the preliminary experiment (**Figure 2B**). We first determined kinetics of mean IAV titers in explants from trachea, bronchi, and lung from all rhesus macaques considered together (**Figure 3A-C**). Mean IAV infection kinetics increased from 0 to 72 hpi in all 3 explant tissues and reached highest levels at 72 hpi for most strains. For most IAV strains, slower kinetics and peak titers that were 10^2-3^ PFU/explant lower were observed in lungs compared to trachea and bronchi. The mean area under the curve (AUC) was significantly higher for H3N8 and H7N5 strains compared to H1N1 and H5N2 strains in trachea and bronchi and trended higher in the lung, although not significantly (**Figure 3D**). Infection kinetics in explants from a representative rhesus macaque (M15) are shown in detail (**Figure 4**) and parallel strain differences were observed in explants from the mean of all 10 animals. Infection kinetics in individual rhesus macaques over time (**Figure 5A**) also reveal the pattern of lower titers in the lung compared to trachea or bronchi. To verify the histologic integrity of explants, hematoxylin and eosin (H&E) staining of tracheal (**Figure 6A-D**) and bronchial explants (not shown) from 2 randomly selected macaques was performed. Microscopically, rhesus macaque tracheal and bronchial explants appeared viable for 48-72 hpi. Normal architecture of ciliated columnar respiratory epithelium, underlying lamina propria, submucosa, and cartilage were generally maintained. Conversely, bronchioles and alveoli were less well preserved. After 48-72 hpi, explants exhibited variable but progressive loss of cilia at the apical epithelial surface, scattered vacuolar degeneration or single-cell necrosis of respiratory epithelial cells, and occasional small foci of epithelial cell loss, though these lesions were also occasionally observed at 0 or 24 hpi. No explants showed evidence of cytopathic effects after IAV inoculation. Together, these results show that multiple IAV subtypes infect viable respiratory tract explants from rhesus macaques with variable kinetics and peak titers.

**Figure 3:**
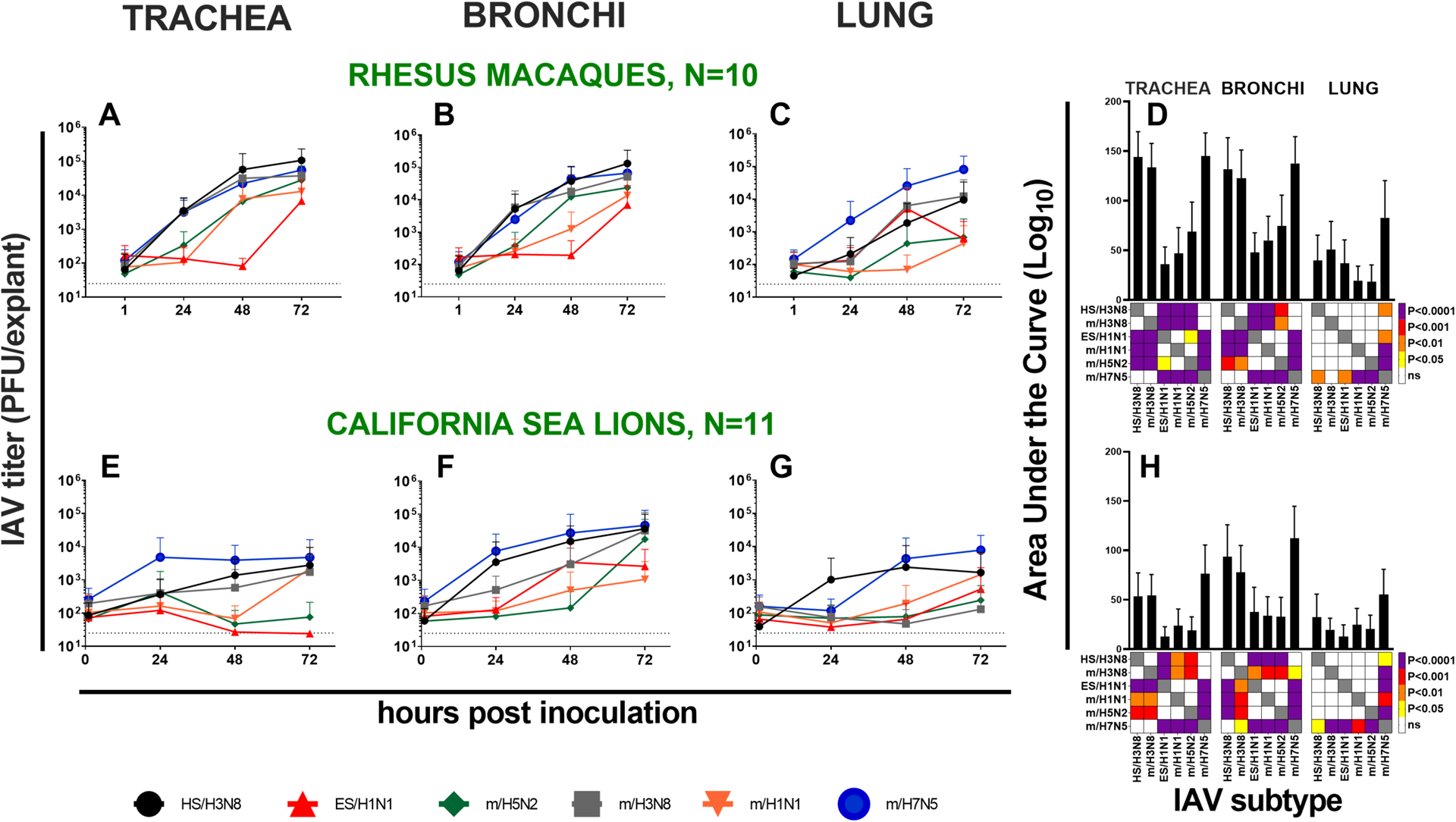
Ex vivo influenza A virus infection kinetics. Mean kinetics in 10 rhesus macaque (**A-C**) and 11 California sea lions (**E-G**) and areas under the infection curve (AUC) (**D, H**) in *ex vivo* respiratory tract trachea, bronchi, and lung explants. Error bars show standard deviations. The dotted black line shows the limit of detection, 50 PFU/explant. Colors in squares in D and H show differences in mean AUC by strain analyzed using one-way ANOVA tests where the darker the color, the smaller the p-value. Strain names refer to the original source of the virus isolate followed by the hemagglutinin and neuraminidase subtype. HS is harbor seal. m is mallard. ES is elephant seal.

**Figure 4.**
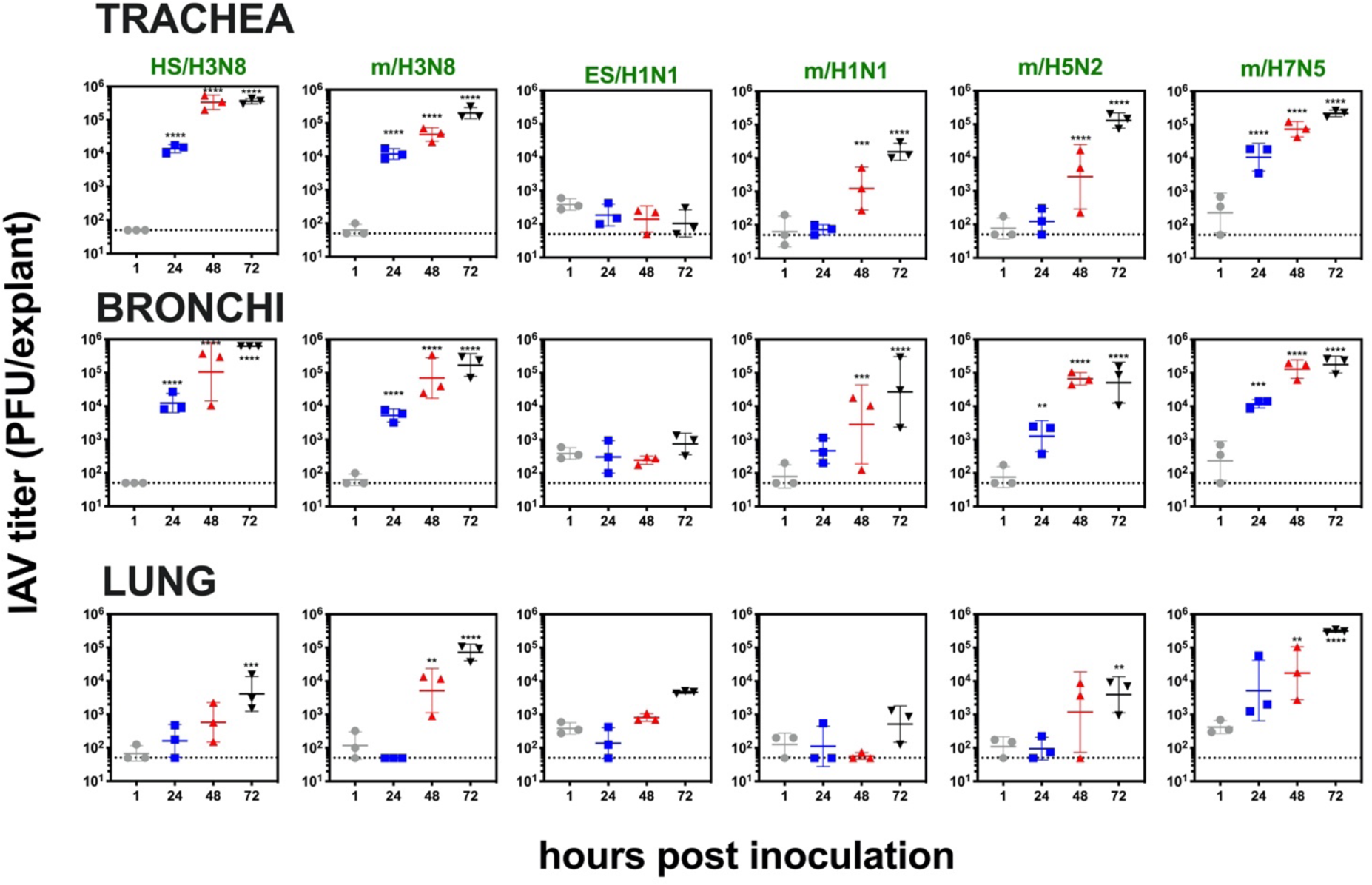
Influenza A virus infection kinetics from 1 to 72 hpi in respiratory tract explants from a 4 year old male rhesus macaque (M15 in Table 2) inoculated with 1 × 10^4^ PFU of 6 viral strains. Viral titers are represented as the geometric mean and geometric standard deviation. Three explants at each time point were titrated independently and the mean of the triplicates is represented by the middle horizontal line. Asterisks show p values (two-way ANOVA with Dunnett’s multiple comparisons) comparing titers at 24, 48 or 72 hpi to 1 hpi, respectively. **p ≤ 0.005, ***p < 0.001, and ****p < 0.0001. The dashed line indicates limit of detection, 50 PFU/explant. Strain names refer to the original source of the virus isolate followed by the hemagglutinin and neuraminidase subtype. HS is harbor seal. m is mallard. ES is elephant seal.

**Figure 5.**
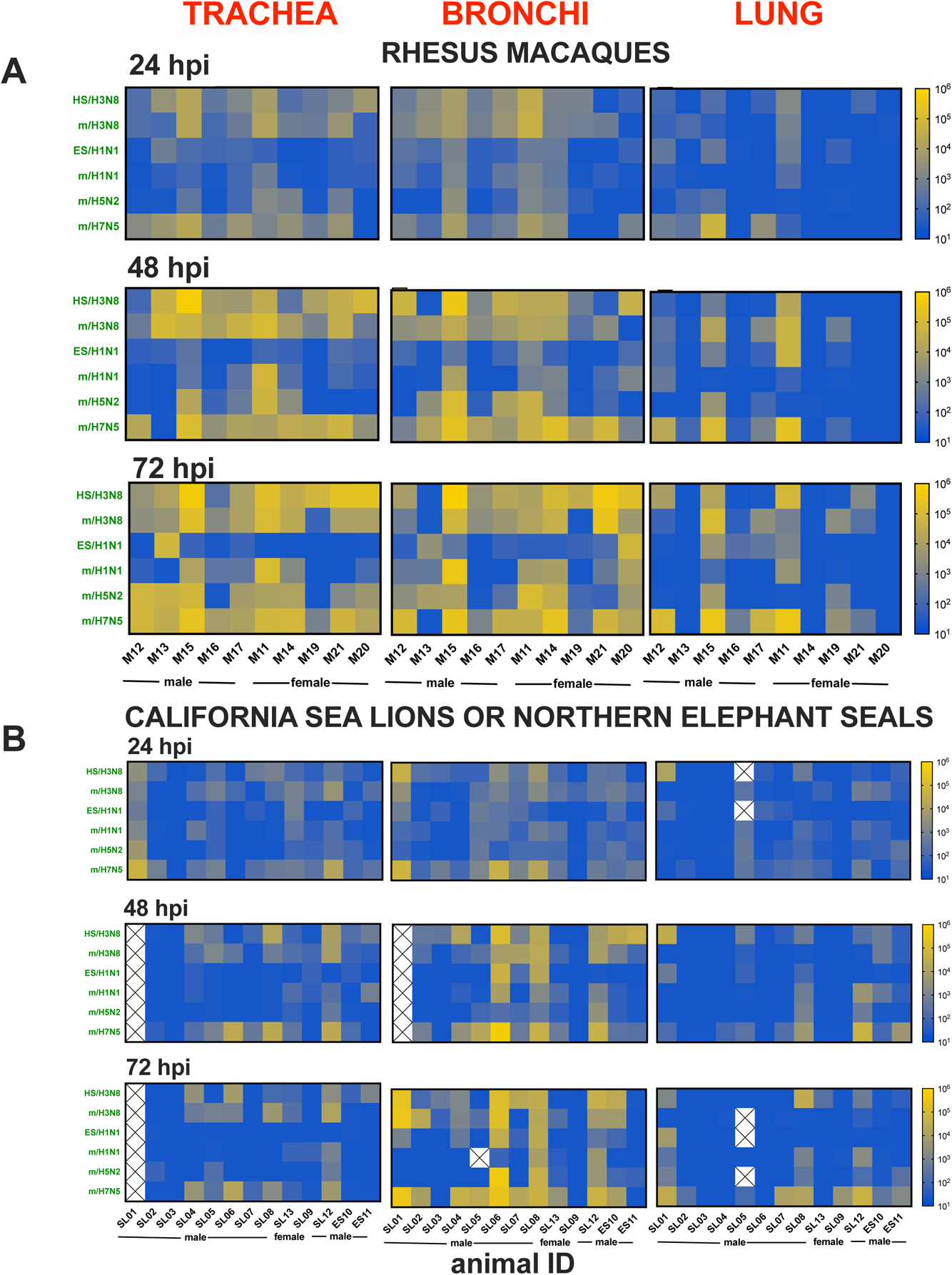
Comparison of *ex vivo* influenza A virus infection kinetics in respiratory tract explants from individual animals. **(A)** IAV titers in explants from 10 rhesus macaques and **(B)** 11 California sea lions and 2 Northern elephant seals inoculated with 1 × 10^4^ PFU and 2 × 10^4^ PFU of each of the 6 IAV strains over a 72 hour period. The kinetics of viral infection were determined by plaque assays of homogenized explants, where each square represents the titer from 10^1^ (blue) to 10^6^ (yellow) in PFU/explant. The limit of detection was 50 PFU/explant. Each explant was titrated once at 2-3 dilutions. Strain names refer to the original source of the virus isolate followed by the hemagglutinin and neuraminidase subtype (Table 1). HS is harbor seal. m is mallard. ES is elephant seal. Boxes with ‘X’ indicate the sample was not available for titration.

**Figure 6.**
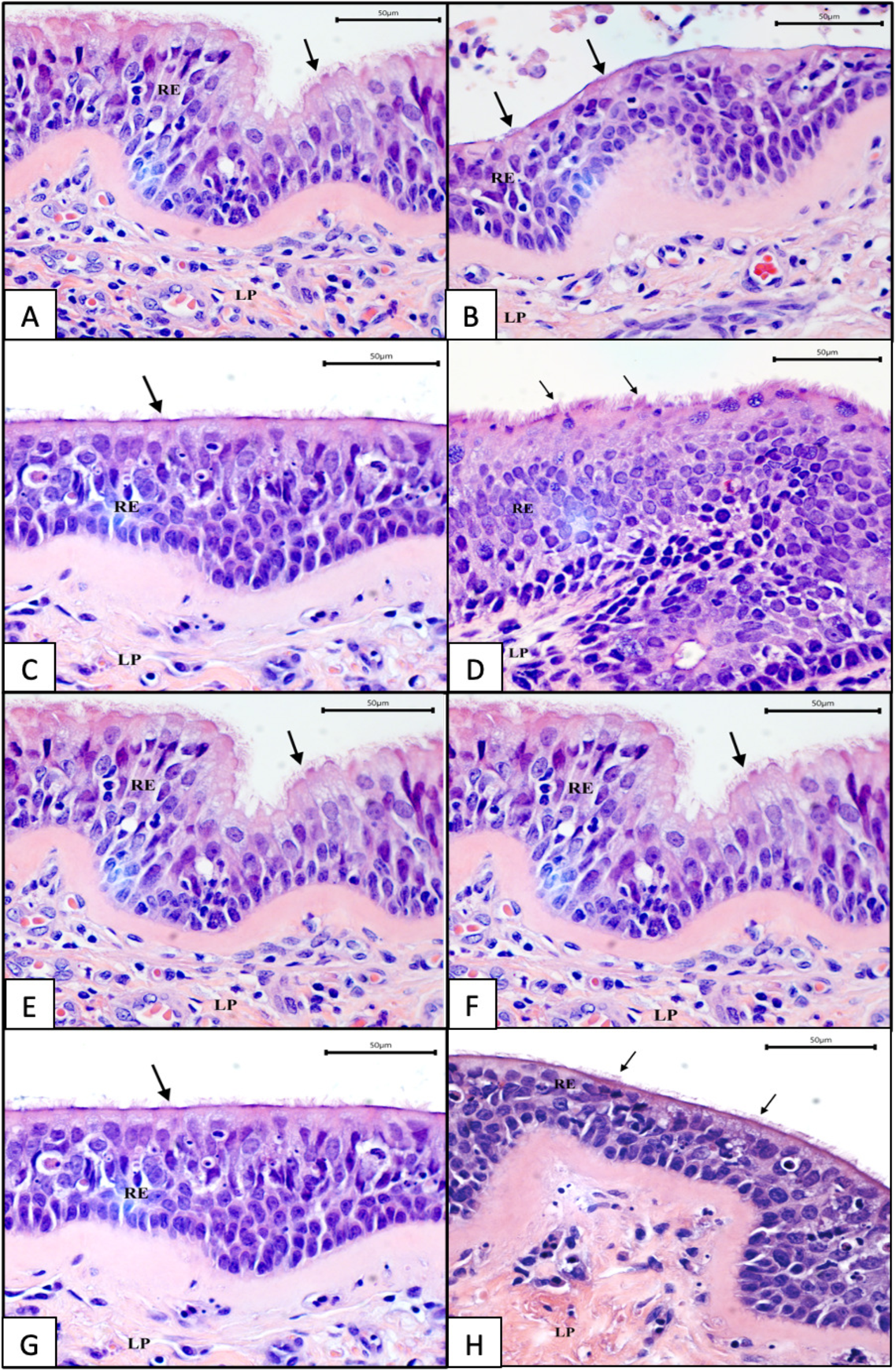
**Representative light photomicrograph of sections of (A-D) rhesus macaque M11 and (E-H) California sea lion SL08 *ex vivo* tracheal explants** at 0 (A, E), 24 (B, F), 48 (C, G), and 72 (D, H) hours. Cilia (black arrows) are present on the apical surface of epithelial cells. Sections were stained with hematoxylin and eosin (400x magnification); bar=50um. RE= respiratory epithelium; LP= lamina propria.

#### Marine mammals

We next examined whether respiratory tract explants from marine mammals are susceptible to infection with the same IAV strains used to infect rhesus macaques, including the 2 strains of marine mammal origin. Explant tissues were obtained from wild California sea lions (N=11) and Northern elephant seals (N=2) euthanized due to non-respiratory medical conditions at The Marine Mammal Center in Sausalito, CA. Marine mammals were mostly male and ranged in age from weanling to adult (**Table 2**). All animals were recovered stranded on beaches. The causes of death varied and were sometimes ascribed to an infectious or toxic etiology but were not typically due to respiratory disease. To assess whether Tissues from animals with gross respiratory tract pathologies at necropsy were also excluded from this study. All California sea lions had a mild burden of a lung worm, *Parafilaroides decorus*, which is found as a normal occurrence in healthy pinnipeds (37). Explants were sectioned to avoid worms and no worms were observed grossly during the IAV infection studies. All marine mammals tested IAV seronegative by ELISA and IAV RNA negative in nasal and rectal swabs at intake to The Marine Mammal Center by RT-PCR (**Table 2**). Although tissues were used within 24 h post-mortem, we confirmed ciliary viability by microscopically visualizing movement of polystyrene beads floating atop cultured California sea lion tracheal tissue from 0 to 72 h. Viability was evidenced by clearance of beads to the side of explants within 60 minutes. As in rhesus macaques, IAV infection kinetics for most strains increased in California sea lions from 0 to 72 hpi in all 3 tissues and reached highest levels at 72 hpi for most strains (**Figure 3E-G**). Similar to rhesus macaques, the mean area under the curve (AUC) was significantly higher for H3N8 and H7N5 strains compared to H1N1 and H5N2 strains in trachea and bronchi while the AUC in lung explants from sea lions was only higher for H7N5 (**Figure 3H**). Detailed data (**Figure 7**) from 1 representative California sea lion (SL12) paralleled patterns from the mean of all 11 animals. Infection kinetics in individual marine mammals over time (**Figure 5B**) showed higher titers in the bronchi relative to trachea or lung. Contrary to our expectation that IAV of marine mammal origin would produce high titers in marine mammal explants, the strain from an elephant seal (ES/H1N1) produced lower and slower kinetics (**Figure 3E-G**, **5B**) than many of the strains isolated from mallards. Like rhesus macaque explants, California sea lion tracheobronchial explants maintained a relatively normal microscopic appearance for 48-72 hours (**Figure 6E-H**), with poor preservation of bronchioles and alveoli and progressive loss of normal architecture over time. All California sea lion tissues also exhibited variable degrees of pre-existing tracheobronchitis, likely associated with their nematode burden. Although California sea lions with respiratory conditions or gross respiratory pathologies were excluded from this study, we also analyzed whether systemic conditions including domoic acid toxicity or septicemia may have exacerbated IAV susceptibility. We assessed AUC for IAV infection kinetics in sea lions euthanized for non-systemic and non-respiratory conditions including malnourishment or leptospirosis (N=6, SL01,SL03-07) versus systemic conditions including domoic acid toxicity or septicemia (N=5, SL02, SL08-13). The AUC infection kinetics between the 2 groups of animals were not significantly different in any of the three respiratory tissues (data not shown). These analyses suggest that IAV infection kinetics in wild California sea lion explants are not impacted by systemic conditions. Together, these results show that, similar to rhesus macaques, multiple IAV subtypes infect viable respiratory tract explants from marine mammals with variable kinetics and peak titers. To further determine whether the differences in mean IAV kinetics were supported statistically over time, we examined IAV infection dynamics in 24 hour windows.

**Figure 7.**
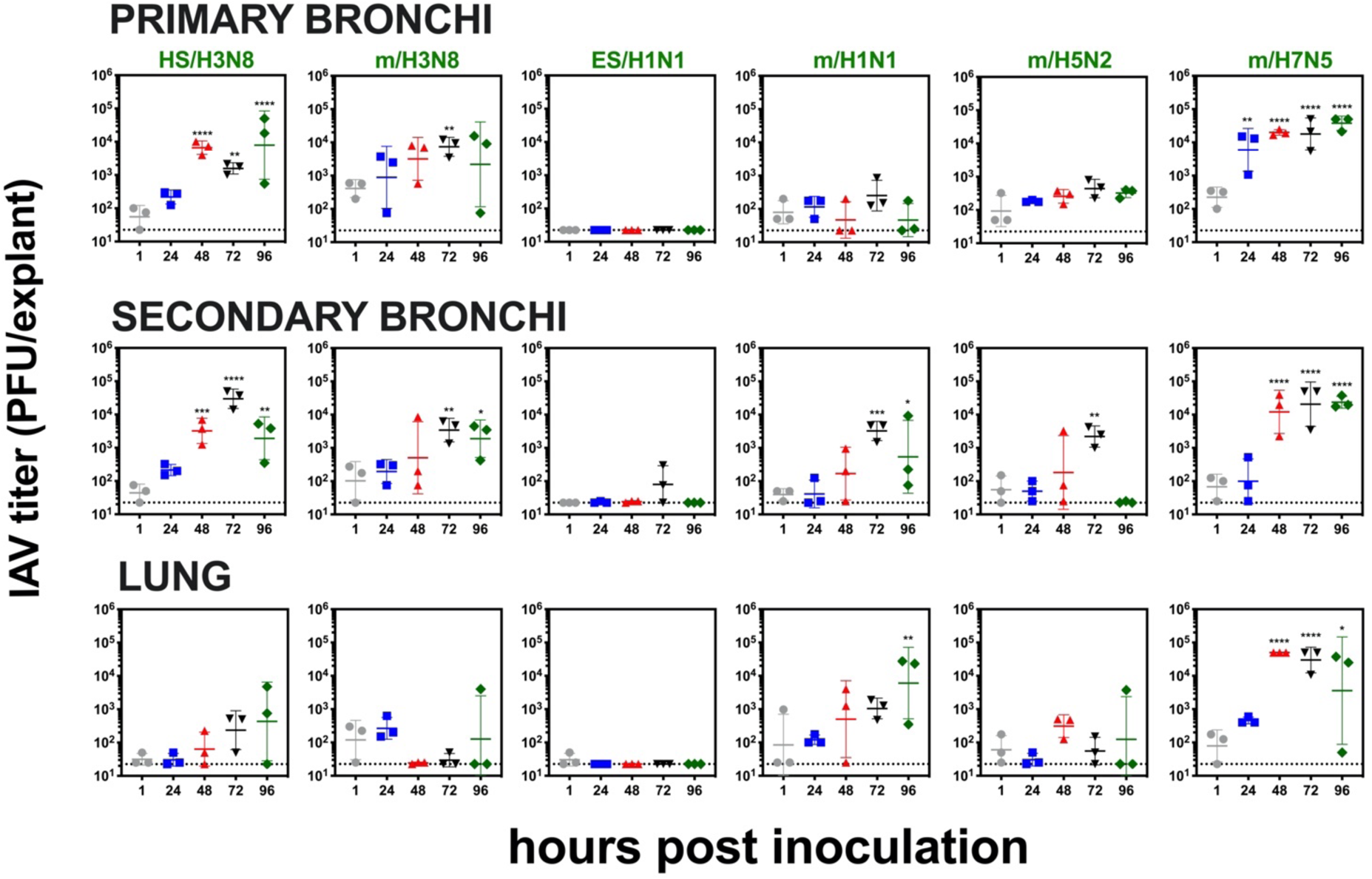
**Influenza A virus infection kinetics from 1 to 96 hpi in respiratory tract explants from an adult female California sea lion** (SL12 in Table 2) inoculated with 2X10^4^ PFU of 6 IAV strains. Viral titers are represented as the geometric mean and geometric standard deviation. Three explants at each time point were titrated independently and the mean of the triplicates is represented by the middle horizontal line. Asterisks show p values (two-way ANOVA with Dunnett’s multiple comparisons) comparing titers at 24, 48, 72 or 96 hpi to 1 hpi, respectively. **p < 0.005, ***p < 0.001, and ****p < 0.0001. The dashed line indicates limit of detection, 50 PFU/explant. Strain names refer to the original source of the virus isolate followed by the hemagglutinin and neuraminidase subtype. HS is harbor seal. m is mallard. ES is elephant seal.

##### IAV Kinetics Over Time

We tested whether explants inoculated with different IAV strains experience different mean changes in titer from 1 to 72 hpi. Considering all 3 tissue types together, the log_10_ change in viral titer was calculated for three time frames: 1 versus 24, 1 versus 48, and 1 versus 72 hpi for rhesus macaques and California sea lions (**Table 3**). Northern elephant seals were not included in analyses since explants from only two animals were available, although they produced infectious IAV after inoculation with some strains (**Figure 5B**). Infection kinetics of the six IAV strains differed for each time frame considered. In rhesus macaques, mean titer changes for HS/H3N8, m/H3N8, and m/H7N5 did not differ significantly from each other over any of the time frames. m/H1N1 and mH5/N2 did not differ significantly from each other over any of the time frames but were different from HS/H3N8, m/H3N8 and m/H7N5. By contrast, the change in ES/H1N1 viral titers significantly differed from all other strains except m/H1N1 from 1 to 48 hpi. In California sea lions, HS/H3N8 and m/H7N5 shared similar mean changes in log_10_ titers in all time frames. For m/H1N1 and HS/H3N8, the mean change in log_10_ titers did not vary significantly from 1 to 24 hpi but did vary from 1 to 48 and 1 to 72 hpi. The mean change in log_10_ viral titers between m/H1N1 and m/H7N5 were only different from 1 to 48 hpi and m/H5N2 and m/H7N5 differed from 1 to 48 and 1 to 72 hpi. These results show that mean changes in IAV titers over 24 hour windows in respiratory tract explants from both rhesus macaques and California sea lions vary with IAV strain.

**Table 3.** Mean change in log_10_ titer of influenza virus strains after inoculation in rhesus macaque or California sea lion respiratory tract explant tissues. Means for the same time frame followed by a common letter are not significantly different by the Games-Howell test at the 5% level of significance. Values in parentheses show standard deviations. The symbols indicate that the mean change in titer differs significantly for at least one strain according to Welch’s ANOVA at 5% level of significance for: *1 vs 24 hpi F=15.417, df=(5,91.649), p-value=5.55e-11 for rhesus macaques and F=4.666, df=(5,189.21), p-value=4.85e-04 for California sea lions, the † denotes 1 vs 48 hpi (F=25.823, df=(5,91.569), p-value=3.59e-16) for rhesus macaques and (F=5.784, df=(5,182.59), p-value=5.55e-05) for California sea lions, and the ‡ denotes 72 hpi (F=23.351, df=(5,91.826), p-value=4.58e-15) for rhesus macaques and (F=6.636, df=(5,189.35), p-value=1.02e-05) for California sea lions. Strain names refer to the original source of the virus isolate followed by the hemagglutinin and neuraminidase subtype. HS is harbor seal. m is mallard. ES is elephant seal.

|  | RHESUS MACAQUES |  |  | CALIFORNIA SEA LIONS |  |  |
| --- | --- | --- | --- | --- | --- | --- |
|  | 1 vs 24 hpi * | 1 vs 48 hpi † | 1 vs 72 hpi ‡ | 1 vs 24 hpi * | 1 vs 48 hpi † | 1 vs 72 hpi ‡ |
| **HS/H3N8** | 1.2 (1.1) <sup>a</sup> | 1.8 (1.5) <sup>ab</sup> | 2.2 (1.5) <sup>ad</sup> | 0.4 (0.8) <sup>e</sup> | 0.6 (1.2) <sup>e</sup> | 0.8 (1.4) <sup>e</sup> |
| **m/H3N8** | 0.9 (1.1) <sup>ac</sup> | 1.8 (1.1) <sup>ac</sup> | 0.9 (1.4) <sup>ab</sup> | -0.04 (0.7) <sup>abc</sup> | -0.07 (1.1) <sup>abc</sup> | 0.2 (1.3) <sup>abc</sup> |
| **ES/H1N1** | -0.1 (0.6) <sup>b</sup> | -0.1 (0.8) <sup>e</sup> | 0.1 (0.9) <sup>e</sup> | -0.1 (0.5) <sup>ab</sup> | -0.1 (0.8) <sup>ab</sup> | -0.1 (0.9) <sup>ab</sup> |
| **m/H1N1** | 0.2 (0.7) <sup>b</sup> | 0.3 (1.0) <sup>de</sup> | 0.9 (1.2) <sup>c</sup> | 0.03 (0.6) <sup>bcde</sup> | -0.05 (0.7) <sup>bd</sup> | 0.2 (1.0) <sup>bcd</sup> |
| **m/H5N2** | 0.3 (0.7) <sup>bc</sup> | 1.0 (1.4) <sup>bcd</sup> | 1.5 (1.4) <sup>bcd</sup> | -0.02 (0.5) <sup>ad</sup> | -0.07 (0.6) <sup>ad</sup> | 0.2 (1.0) <sup>ad</sup> |
| **m/H7N5** | 1.1 (0.8) <sup>a</sup> | 1.9 (1.0) <sup>a</sup> | 2.4 (1.2) <sup>a</sup> | 0.3 (0.9) <sup>cde</sup> | 0.5 (1.2) <sup>ce</sup> | 0.8 (1.5) <sup>ce</sup> |

##### IAV Kinetics by Tissue

To examine whether tissue type associated with IAV titer, we analyzed all 6 IAV strains together (**Table 4**). Infection kinetics in bronchi and trachea from rhesus macaques were significantly higher than in the lung over all 24 hour windows from 1 to 72 hpi. In California sea lions, infection kinetics in bronchi and trachea were significantly higher than in the lung at 24 hpi, while infection kinetics in bronchi were significantly higher than in the trachea and lung at 48 and 72 hpi. Together, these data support tissue-specific IAV infectivity, where bronchi and trachea from both species support production of higher IAV titers than lung explants.

**Table 4.** Mean change in log_10_ titer of influenza virus in respiratory tract tissues from rhesus macaques and California sea lions. Means for the same time frame followed by a common letter are not significantly different by the Games-Howell test at the 5% level of significance. Values in parentheses show standard deviations. The symbols indicate that the mean change in titer differs significantly for at least one tissue according to Welch’s ANOVA at 5% level of significance for: * 1 vs 24 hpi (F=28.071, df=(2, 127.39), p-value=7.93e-11) in rhesus macaques, (F=15.369, df=(2, 269.45), p-value=5.96e-05) for California sea lions, † 1 vs 48 hpi (F=13.641, df=(2, 129.47), p-value=4.02e-6) in rhesus macaques, (F=10.105, df=(2, 257.71), p-value=5.96e-05) in California sea lions, ‡ 1 vs 72 hpi (F=7.8203, df=(2, 133.05), p-value=6.16e-4) for rhesus macaques, (F=11.083, df=(2, 268.89), p-value=2.37e-0.5) for California sea lions.

|  | RHESUS MACAQUES |  |  | CALIFORNIA SEA LIONS |  |  |
| --- | --- | --- | --- | --- | --- | --- |
| Tissue | 1 vs 24 hpi<sup>*</sup> | 1 vs 48 hpi<sup>†</sup> | 1 vs 72 hpi<sup>‡</sup> | 1 vs 24 hpi<sup>*</sup> | 1 vs 48 hpi<sup>†</sup> | 1 vs 72 hpi<sup>‡</sup> |
| Bronchi | 1.0 (0.9)<sup>a</sup> | 1.5 (1.4)<sup>a</sup> | 1.7 (1.5)<sup>a</sup> | 0.3 (0.8)<sup>a</sup> | 0.5 (1.3)<sup>a</sup> | 0.8 (1.5)<sup>a</sup> |
| Trachea | 0.8 (1.0)<sup>a</sup> | 1.4 (1.5)<sup>a</sup> | 1.9 (1.5)<sup>a</sup> | 0.1 (0.7)<sup>a</sup> | 0.1 (0.8)<sup>b</sup> | 0.2 (1.0)<sup>b</sup> |
| Lung | 0.1 (0.7)<sup>b</sup> | 0.5 (1.1)<sup>b</sup> | 1.0 (1.4)<sup>b</sup> | -0.2 (0.6)<sup>b</sup> | -0.1 (0.8)<sup>b</sup> | 0.1 (1.1)<sup>b</sup> |

##### IAV Kinetics by Species

To evaluate whether species associates with infection kinetics for the 6 IAV strains, we compared changes in mean log_10_ titer between rhesus macaque and California sea lion explants (**Table 5**). The mean change in titer of both H3N8 strains and m/H7N5 was significantly higher in trachea at all time points and in bronchi to 48 hpi in rhesus macaques compared to California sea lions. Changes in mean log_10_ titers for m/H3N8 in the lung from 1 to 48 and 1 to 72 hpi were higher in rhesus macaques compared to California sea lions. Together these data show that rhesus macaque explants produce higher titers compared to California sea lions for most IAV strains used here.

**Table 5.** Difference in mean log_10_ change for respiratory tract explant tissues from California sea lions versus rhesus macaques. Differences are show as p-values after the Games-Howell post-hoc test. Significant *p*-values (*p* < 0.05) are in bold. Strain names refer to the original source of the virus isolate followed by the hemagglutinin and neuraminidase subtype. HS is harbor seal. m is mallard. ES is elephant seal. hpi is hours post inoculation.

| Timeframe | IAV Strain | Trachea | Bronchi | Lung |
| --- | --- | --- | --- | --- |
| 1-24 hpi | HS/H3N8 | **-1.26 (0.011)** | **-1.13 (0.023)** | -0.11 (0.989) |
|  | m/H3N8 | **-1.26 (0.003)** | **-1.40 (<0.001)** | -0.29 (0.695) |
|  | ES/H1N1 | -0.03 (1.00) | -0.13 (0.967) | -0.10 (0.996) |
|  | m/H1N1 | -0.14 (0.959) | -0.32 (0.475) | -0.03 (1.00) |
|  | m/H5N2 | -0.41 (0.362) | -0.45 (0.388) | -0.07 (0.997) |
|  | m/H7N5 | **-1.02 (0.047)** | -0.62 (0.266) | -0.46 (0.551) |
| 1-48 hpi | HS/H3N8 | **-1.95 (0.002)** | -1.27 (0.120) | -0.18 (0.982) |
|  | m/H3N8 | **-2.21 (<0.001)** | **-1.50 (0.009)** | **-1.50 (0.006)** |
|  | ES/H1N1 | -0.02 (1.00) | 0.25 (0.963) | -0.45 (0.572) |
|  | m/H1N1 | -0.74 (0.160) | -0.48 (0.714) | 0.01 (1.00) |
|  | m/H5N2 | -0.99 (0.100) | **-1.50 (0.024)** | -0.25 (0.918) |
|  | m/H7N5 | **-1.75 (<0.001)** | **-1.50 (0.001)** | -0.71 (0.540) |
| 1-72 hpi | HS/H3N8 | **-2.41 (<0.001)** | **-1.74 (0.015)** | -0.49 (0.711) |
|  | m/H3N8 | **-2.02 (<0.001)** | -1.44 (0.085) | **-1.87 (0.006)** |
|  | ES/H1N1 | -0.05 (1.00) | 0.01 (1.00) | -0.48 (0.464) |
|  | m/H1N1 | -0.83 (0.265) | -1.06 (0.131) | -0.03 (1.00) |
|  | m/H5N2 | **-2.11 (<0.001)** | -1.23 (0.083) | -0.32 (0.893) |
|  | m/H7N5 | **-2.16 (<0.001)** | -1.08 (0.229) | -1.11 (0.335) |

### Cellular tropism of influenza A virus in rhesus macaque and California sea lion explants

We used immunohistochemistry to assess the cellular tropism of IAV in rhesus macaque and California sea lion respiratory tissue explants (**Figure 8**, **Table 6**). At least 22 sections from each tissue type for 2 rhesus macaques (M11 and M13) and at least 56 sections from each tissue type for 2 California sea lions (SL08 and SL12) were evaluated. All examined sections exhibited variable degrees of nonspecific cytoplasmic, membranous and/or background staining; as such, only cells with strong nuclear immunoreactivity were considered positive. For all explants from both species and all IAV subtypes, positive staining for viral nucleoprotein was limited to epithelial cells of the trachea and bronchi; representative slides are shown (**Figure 8A-D**). Staining for IAV nucleoprotein was not observed in non-inoculated explants or IHC antibody-treated negative isotype controls from rhesus macaque (**Figure 8E-F**) or California sea lion (**Figure 8G-H**) trachea. Pneumocytes within the lung were IAV negative. Infection was primarily localized to apical epithelial cells, particularly tracheal and bronchial ciliated columnar cells and goblet cells, while basal cells were largely spared. Rhesus macaque explants from M11 and M13 infected with H3N8 strains exhibited higher relative proportions of positive tracheal and bronchial epithelial cells 24 and 48 hpi, even though IAV titers at those times for most strains exceeded 10^4^ PFU/explant 24 hpi (**Figure 5A**). The explants from M11 infected with m/H7N5 exhibited the largest variation in relative proportion of positive cells between 24 and 48 hpi. The number of infected respiratory epithelial cells generally increased between 24 and 48 hpi, which is consistent with infection kinetics where IAV titers increased during that period. While positive nuclei were less frequent within California sea lion explants, those infected with H3N8 strains also exhibited higher relative proportions of positive tracheal and bronchial epithelial cells at 24 and 48 hpi, which is consistent with data from rhesus macaques. These results demonstrate that IAV infects the apical epithelial cells of upper respiratory tract explants in rhesus macaques and California sea lions.

**Figure 8.**
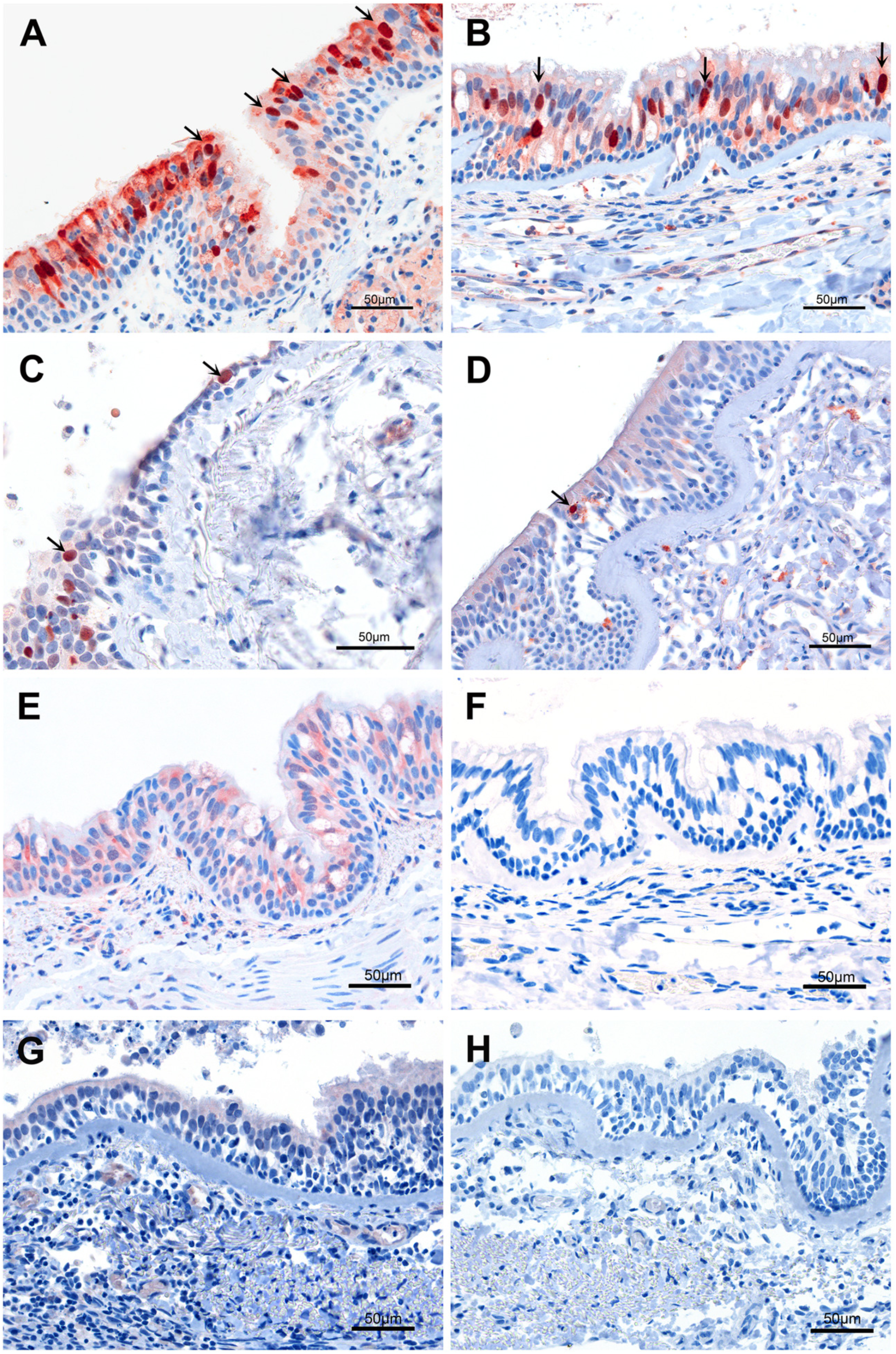
Immunohistochemical (IHC) staining for influenza A virus in rhesus macaque and California sea lion tracheal and bronchial explants. **(A)** Bronchi from rhesus macaques 48 hours post inoculation (hpi) with **(A)** m/H7N5 (animal M11) or **(B)** HS/H3N8 (animal M13); **(C)** Trachea from California sea lion SL12 48 hpi with HS/H3N8; **(D)** Trachea from California sea lion SL08 24 hpi with m/H3N8; Bronchi from rhesus macaque M13, 48 hpi treated with **(E)** growth medium only or **(F)** with HS/H3N8 and stained with isotype IgG; Trachea from SL12 24 hpi treated with **(G)** growth medium only or **(H)** with HS/H3N8 and stained with isotype IgG. IHC stain used an antibody that labels IAV nucleoprotein. Positive respiratory epithelial cells exhibit strong nuclear immunoreactivity (red-brown nuclear staining; arrows). Positive staining is primarily localized to apical epithelial cells, with relative sparing of basal cells.

**Table 6.**
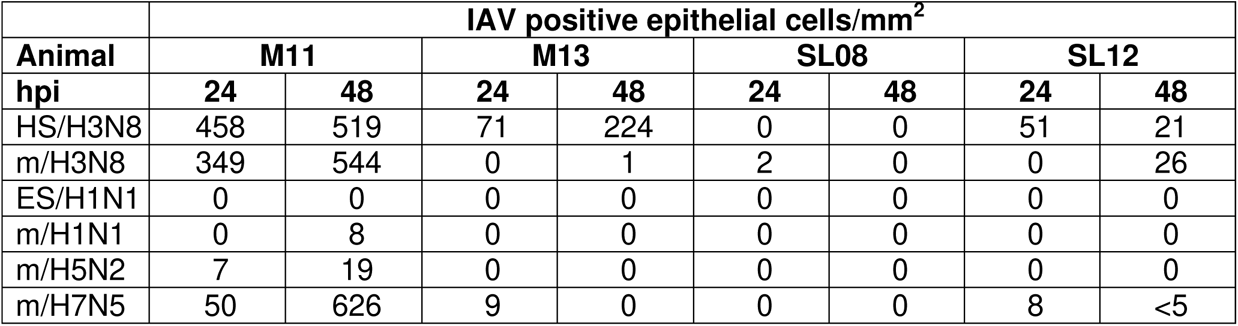
Immunohistochemical detection of influenza A virus in macaque and California sea lion respiratory tissue explants. Trachea, bronchi and lung were examined for each animal. At least 22 sections from each tissue type were examined for the rhesus macaques and at least 56 sections from each tissue type were examined for the California sea lions. Strain names refer to the original source of the virus isolate followed by the hemagglutinin and neuraminidase subtype. HS is harbor seal. m is mallard. ES is elephant seal. hpi is hours post inoculation.

### Genetic changes in influenza A viruses sequenced from explants

IAV genomes in inoculated explants from 2 rhesus macaques, 3 California sea lions, and 1 Northern elephant seal were compared to sequences of the strains used as inocula (**Table 7**). Sequence comparisons showed that all genomes from rhesus macaque explants were 100% identical at the consensus level (mutations occurring on 50% or more of viral RNAs) to their corresponding inoculum (data not shown). Six nonsynonymous nucleotide substitutions in the HA, NA, and PA genes were detected in California sea lion and Northern elephant seal explants inoculated with different IAV subtypes (**Table 8**). Inocula contained a mixture of bases including the mutant nucleotide at each of these loci, indicating that none of the mutations developed *de novo* in explants (data not shown). One synonymous nucleotide substitution, PB2 1368 in m/H7N5, was detected in bronchi from all 3 California sea lions at 48 and 72 hpi. The nucleotide affecting this substitution was present at a minority frequency (41%) in the inoculum, indicating that it did not develop *de novo* but increased to consensus frequency in explants by 48 hpi (data not shown). Together these data show that no *de novo* IAV mutations were detected in any strain during *ex vivo* respiratory tract explant infection from selected rhesus macaques and marine mammals.

**Table 7.** IAV inoculated rhesus macaque and marine mammal respiratory tract explants from which whole genome IAV sequences were obtained at indicated times post-inoculation and compared to their respective inocula. Empty squares indicate samples for which sequencing was not attempted. HS is harbor seal. m is mallard. ES is elephant seal, M is macaque, SL is California sea lion.

|  |  | hour(s) post inoculation |  |  |  |  |  |
| --- | --- | --- | --- | --- | --- | --- | --- |
| **Animal ID** | **tissue** | **HS/H3N8** | **m/H3N8** | **ES/H1N1** | **m/H1N1** | **m/H5N2** | **m/H7N5** |
| M15 | trachea | 72 | 72 |  | 72 | 72 | 72 |
|  | lung |  |  |  |  |  | 72 |
| M11 | trachea | 72 | 72 |  | 72 |  | 72 |
| SL01 | bronchi |  | 72 |  |  |  | 72, 96 |
| SL06 | bronchi | 48, 72 | 72 |  |  | 72 | 48, 72 |
| SL08 | trachea |  |  |  |  |  | 48 |
|  | bronchi |  |  | 72 |  |  | 72 |
| ES10 | bronchi | 48, 72 |  |  |  |  |  |

**Table 8.** IAV amino acid changes compared to inocula detected in ex vivo bronchi explants from California sea lions and a Northern elephant seal. Numbers shown correspond to positions in indicated IAV proteins. *Indicates that the mutation was a synonymous change. Empty squares indicate no sequence differences were detected. An X shows samples for which sequencing was not attempted. hpi is hours post inoculation, HS is harbor seal, m is mallard, ES is elephant seal, M is macaque, SL is California sea lion, HA is hemagglutinin, NA is neuraminidase, PA is polymerase acidic protein, PB is polymerase basic protein.

| Animal ID | hpi | HS/H3N8<br>HA | HS/H3N8<br>NA | ES/H1N1<br>PA | m/H7N5<br>HA | m/H7N5<br>PB2* |
| --- | --- | --- | --- | --- | --- | --- |
| SL01 | 72 |  |  |  | V311I | 1368 |
|  | 96 |  | V394I |  |  |  |
| SL06 | 48 |  |  |  |  | 1368 |
|  | 72 |  | P118S |  |  | 1368 |
| SL08 | 48 |  |  |  |  | 1368 |
|  | 72 |  |  | I359N,<br>M441I |  | 1368 |
| ES10 | 48 | A154T | P118S |  |  |  |
|  | 72 | A154T | P118S |  |  |  |

## Discussion

We adapted an *ex vivo* respiratory tract infection model to study susceptibility and to define comparative infection kinetics of IAV in colony-raised rhesus macaques and wild marine mammals. We observed that IAV exhibits temporal, subtype- and species-dependent infection kinetics in explants from both rhesus macaques and California sea lions. Although the relative infection kinetics for the six strains was similar for explants from rhesus macaques and California sea lions, similar patterns were not observed in MDCK cells. This suggests that immortalized cell lines may not accurately represent infection phenotypes *in vivo*, further underscoring the value of using *ex vivo* systems.

E*x vivo* respiratory tract models have been used for studying respiratory pathogens of humans, swine, bovines, canines, and equines (23, 25–28, 38–40). *Ex vivo* cultures of respiratory tissues provide a close resemblance to the respiratory tract *in vivo* by: 1) shared polarity, where the basolateral surface is exposed to culture medium and the apical cell surface is exposed to air, 2) shared cell positioning *in situ* which maintains virus receptor distribution, 3) possession of multiple cell types and states of differentiation, 4) maintenance of the three dimensional integrity and architecture of a tissue, unlike cell monocultures, and 5) ciliary activity to preserve mucociliary clearance. Given these similarities to living IAV hosts, explants are valuable tools for assessing host susceptibility, infection kinetics, and pathogenesis of respiratory pathogens. *Ex vivo* systems also better assess infectivity compared to cell monoculture binding assays that only measure virus-receptor affinity (25, 27). In addition, tissues collected from a single animal can be divided into pieces and used to compare relative infectivity of different viral strains while holding the host constant. Last, use of explants from animals euthanized for other reasons reduced the number of animals used in research, following the principles of reduction, replacement, and refinement.

Limitations of the *ex vivo* approach include restricted availability of tissues, short periods of viability, and inter-animal variability in IAV susceptibility. The absence of a blood supply also constitutes a weakness since recruitment of immune cells to the infection site cannot occur. However, such a pitfall might be co-opted in future studies to understand the effects of innate and intrinsic immunity without immune cell influx. Use of explants from wild marine mammals euthanized for non-respiratory conditions, some of which present systemically, is another limitation of this study. Some of the marine mammals used in this study showed evidence of exposure to toxins, malnourishment, or congenital defects, factors that may modify IAV susceptibility. However, systemic conditions including toxin exposure did not correspond to observed differences in IAV infection capacity in California sea lion explants in the current study. Given that the only marine mammals available for this study were those euthanized for other conditions, we unfortunately do not have the opportunity to study tissues from healthy animals, especially since both California sea lions and elephant seals are protected by the Marine Mammal Protection Act.

Lower IAV titers in *ex vivo rhesus* macaque and marine mammal lungs relative to trachea and bronchi support the upper respiratory tract as the primary site of virus infection with the virus strains used in this study. The ability of IAV to produce higher titers in the upper respiratory tract may reflect adaptation of the virus to infect cell targets in closest anatomic proximity to its entry point and shedding site, the respiratory mucosa. Lower susceptibility of the lung to IAV infection could result from decreased receptor expression in that tissue. We did not identify or enumerate IAV receptor expression in explants in this study to confirm this hypothesis. An alternate possibility is that lungs of both species decayed more quickly than the trachea and bronchi, rendering them less susceptible to IAV infection. However, we feel accelerated lung decay is unlikely given our gross and histologic observations showed similar architectural integrity in all 3 tissues to 72 hpi (data not shown). Alternately, lower IAV lung infectivity could result from increased resident immune cell activation and response in that tissue.

The infection kinetics of harbor seal-origin H3N8 and mallard-origin H7N5 in California sea lion respiratory tract explant tissues were superior to other subtype strains isolated from mallards and elephant seal-origin H1N1. Since all strains share similar passage histories, infection kinetic differences are not likely due to varying passage or changes in consensus sequence since all were nearly identical to those of their unpassaged progenitors. Our observation that the elephant seal H1N1 infection kinetics were lower than other strains in California sea lions suggests that having a marine mammal as the isolate source does not necessarily translate to higher marine mammal explant infectivity. The lack of augmented infectivity in explants of the species from which the isolate was made has also been observed in swine, where strains of the same subtype isolated from both swine and humans showed similar infection kinetics (41). Further, some human-isolated IAV show great inter-strain variability in infection kinetics in human explants (27). In spite of the different infection kinetics across strains, the 6 strains used here, representing 4 IAV subtypes, productively infected at least one explant type for both species. These data suggest that any of these subtypes could also infect marine mammals in the wild, consistent with serologic data showing exposure to many subtypes (13–16, 18, 19). Together, data presented here show that the marine mammal *ex vivo* culture of respiratory tissues is a tool to study IAV susceptibility, host-range, and tissue tropism.

## ACKNOWLEDGEMENTS

We are grateful to the veterinary and research staff at The Marine Mammal Center for coordinating the collection and transfer of fresh tissue samples to our laboratory. Sample possession was granted by a Letter of Authorization from the National Oceanic and Atmospheric Administration National Marine Fisheries Service.

## FUNDING ACKNOWLEDGEMENTS

Funding support was provided by NIH NIAID #HHSN272201400008C to WB and LLC and from the University of California, Davis Office of Research to PAP and HL. EEB was supported by a Comparative Medical Science Training Program T32 fellowship # OD011147. We thank Kathy West at the California National Primate Research Center for generating the rhesus macaque image used in figures and Ana L. Ramirez for generating the other images used in figures.

## AUTHOR CONTRIBUTIONS

Conceptualization: HL, MP, WB, LLC, Methodology: HL, CMW, LLC, BH, Investigation: HL, MP, CMW, OG-V, AMA, EEB, ZMM, KMH, KJ, PAP, ZK, DK. Writing-original draft: HL, LLC. Writing-review and editing: HL, CMW, MP, EEB, ZMM, HB, BH, KMH, WB, PAP, LLC. Visualization: HL, CMW, EEB, PAP, KJ, LLC. Supervision and project administration: WB, HB, LLC. Funding acquisition: WB, HL, PAP, LLC.

